# Transferrin receptor 1 (TfR1) functions as an entry receptor for scale drop disease virus to invade host cell via clathrin-mediated endocytosis

**DOI:** 10.1101/2025.04.15.648974

**Authors:** Jiaming Chen, Yuting Fu, Yong Li, Shaoping Weng, Hebing Wang, Jianguo He, Chuanfu Dong

## Abstract

Scale drop disease virus (SDDV), a distinct member of the genus *Megalocytivirus* within the *Iridoviridae* family, has emerged as a novel threat to global teleost aquaculture. Despite its importance, the pathogenic mechanism of SDDV remains largely elusive. In this study, we identified mandarin fish transferrin receptor 1 (*mf*TfR1) as an entry receptor for SDDV to invade host cells. Firstly, *mf*TfR1 was detected in high abundance in purified SDDV virions and exhibited dynamic responses to SDDV infection, showing distinct regulatory patterns both *in vivo* and *in vitro*. Overexpression of *mf*TfR1 in low-permissive fathead minnow (FHM) cells significantly enhanced SDDV replication, particularly during the early stages of viral binding and entry. Conversely, antibody-blocking experiments and treatment with the TfR1 inhibitor ferristatin II significantly suppressed SDDV entry. Further investigation revealed that *mf*TfR1 directly interacted with the major capsid protein (MCP) of SDDV, and the helical domain of *mf*TfR1 was identified as the crucial docking site. The binding site within the helical domain was determined, and disrupting this interaction significantly reduced viral entry and host mortality. Finally, we demonstrated that SDDV could activate Src kinase-mediated tyrosine phosphorylation of *mf*TfR1. This phosphorylation event enhanced the internalization of *mf*TfR1 and facilitated clathrin-mediated endocytosis. Collectively, our study provides compelling evidence to confirm that *mf*TfR1 functions as an entry receptor that mediates SDDV entry into host cells via clathrin-mediated endocytosis, leading to a lethal infection outcome. Our work lays the ground work for the development of targeted therapeutic strategies to mitigate the impact of SDDV in aquaculture.

**IMPORTANCE:** TfR1, a dimeric glycoprotein classified as a type II transmembrane receptor, facilitates the cellular internalization of holo-transferrin. In several mammalian and avian RNA viruses, TfR1 serves as a crucial receptor to mediate the entry of viruses into host cells. As an emerging large DNA virus, SDDV poses an emerging threat to teleosts globally, however, its underlying pathogenic mechanisms remain poorly understood. In this study, we are the first to identify *mf*TfR1 as a crucial receptor for SDDV entry. We demonstrated a specific interaction between *mf*TfR1 and the major capsid protein (MCP) of SDDV, with the helical domain of *mf*TfR1 acting as the binding site. Moreover, we confirmed that SDDV enters cells through *mf*TfR1-mediated clathrin-dependent endocytosis. This work highlights the essential role of TfR1 in aquatic DNA viral infections and establishes the theoretical foundation for developing targeted therapeutic strategies against SDDV.

## INTRODUCTION

Scale drop disease virus (SDDV), a large double-stranded DNA virus with genome size of approximately 131 kbp, is a novel member of the genus *Megalocytivirus* within the family *Iridoviridae* (1-3). As early as the early 1990s, it was associated with severe cases of scale drop syndrome (SDS) of farmed Asian sea bass (*Lates calcarifer*, also known as barramundi) in Penang, Malaysia (3, 4). The causative agent of SDS in cultured *L. calcarifer* was directly observed under a transmission electron microscope (TEM) as an iridovirus-like virus. However, it was further confirmed to be a non-red sea bream iridovirus (RSIV) since it could not be recognized by the anti-RSIV monoclonal antibody (4). Until 2015, the virus was isolated and characterized at cell culture level and identified as a distinctive megalocytivirus, and designated as SDDV (3). So far, SDDV was widely documented in SDS associated *L. calcarifer* from several Southeast Asian countries, including Singapore, Malaysia, and Thailand, and has become a major concern in Asian sea bass breeding in these regions (5-10). The cumulative mortality of SDDV-infected Asian seabass can reach 40∼50%, affecting both juvenile and adult fish (3, 5, 11). In China, SDDV was firstly isolated from the ascitic fluid of yellowfin seabream (*Acanthopagrus latus*) with severe ascites in Zhuhai, Guangdong Province in 2020 (2). The comparative genome investigation showed that SDDV from yellowfin seabream in China was >99% identical to those isolates from Asian seabass in SE Asian country (2, 7), indicating that they have the same origin. The epidemiological investigation of diseased yellowfin seabream in China showed that the positivity rate of SDDV detected by conventional PCR in diseased yellowfin seabream from Zhuhai was 38.24%, with mortalities in affected ponds exceeding 60%, indicating the widespread SDDV transmission and significant loss outcome (12). Additionally, very low doses of SDDV could cause 100% mortality in mandarin fish (*Siniperca chuatsi*), suggesting it as a potential host fish species for SDDV transmission (13). Additional to SDDV in Asian sea bass, yellowfin sea bream and mandarin fish in Asia, SDDV close viruses were also reported in European chub (*Squalius cephalus*) from England (14), and tilapia (*Oreochromis spp*.) from the USA (15).

According to the latest on line version of the International Committee on Taxonomy of Viruses (<u>Genus: Megalocytivirus | ICTV</u>), the genus *Megalocytivirus* is divided into two species, namely *Megalocytivirus lates1* and *M. pagrus1*, respectively, and SDDV and infectious spleen and kidney necrosis virus (ISKNV)/RSIV are considered the type isolates of these two species. Compared with ISKNV/RSIV, which has been extensively studied (16-19), study on SDDV still remains in preliminary stage. Using LC-MS/MS, we recently identified 65 viral proteins from purified SDDV virions, including two envelope proteins (VP056 and VP098) and one capsid protein (VP033) (12). Comparative analysis of the structural proteins between SDDV and ISKNV/RSIV revealed substantial differences, suggesting that SDDV is a unique species within genus *Megalocytivirus*. Extensive research has been conducted on SDDV both domestically and internationally, focusing primarily on epidemiological investigations, diagnostic techniques (20), as well as vaccine development (2, 3, 10, 21). However, studies underlying the pathogenic mechanisms of SDDV remain scarce. Our recent study revealed that SDDV infection but not ISKNV or mandarin fish ranavirus (MRV) infection could induce classical ferroptosis both *in vitro* mandarin fish fry cell line and *in vivo* mandarin fish model (22), and ferroptosis was identified as the primary mode of cell death upon SDDV infection. Further investigation demonstrated that mandarin fish transferrin receptor-1 (*mf*TfR1) plays a critical role in SDDV-induced ferroptosis, as downregulation of *mf*TfR1 significantly reduces SDDV replication and consequently decreases the mortality in infected mandarin fish (22), however, the exact role of *mf*TfR1 involvement in infection and pathogenicity of SDDV remain unknown.

TfR1, a dimeric glycoprotein categorized as a type II transmembrane receptor, facilitates the cellular internalization of holo-transferrin. This process occurs via clathrin-mediated endocytosis (CME), thereby enabling the delivery of iron (23). Generally, its expression is tightly controlled by intracellular iron concentrations. Disruption of iron regulatory mechanisms under specific pathological conditions can activate ferroptosis, a form of iron-dependent cell death (24-26). TfR1 can translocate from the Golgi apparatus to the plasma membrane, where it forms a complex with holo-transferrin. This interaction facilitates the internalization of extracellular iron into the cellular machinery, ultimately triggering ferroptosis (27, 28). Beyond its canonical roles in iron acquisition and intestinal epithelial homeostasis (11, 29), TfR1 serves as a critical host factor for viral entry. Notably, it facilitates infection of New World arenaviruses (e.g., Machupo virus) (30, 31), mouse mammary tumor virus (MMTV) (32-34), canine and feline parvoviruses, feline panleukopenia virus (35, 36), poliovirus (37) and hepatitis C virus (HCV) (38). Moreover, Junín virus (JUNV) utilizes TfR1 as a cellular receptor to enter susceptible cells via CME (39, 40). While rabies virus and SARS-CoV-2 require cooperative interactions between TfR1 and mGluR2 to internalize viral glycoproteins via clathrin-coated vesicles (41). Similarly, TfR1 engages the spike protein of porcine epidemic diarrhea virus (PEDV) to initiate infection (42). In summary, TfR1 interacts with specific viral proteins, thereby playing a pivotal role during virus infections (34, 42). Although its functions have been extensively investigated in mammals and avians, research on fish TfR1, especially its role in teleost anti-viral responses, remains scarce. Within the field of virology, the majority of studies have been centered on RNA viruses. In contrast, research on large DNA viruses, like scale drop disease virus (SDDV), has made little headway. This is primarily due to their larger genomes and the more complex nature of their viral proteins. These characteristics pose significant challenges to traditional research methods and hamper our understanding of their biological mechanisms.

Our previous studies identified spleen, an important organ for iron storage, as the primary target tissue of SDDV infection (2, 21). Moreover, we found that SDDV infection induces ferroptosis in mandarin fish, which is closely linked to *mf*TfR1 (22). In this study, we employed mandarin fish as an *in vivo* model and mandarin fish fry (MFF-1) cells for *in vitro* experiments to characterize the interaction between *mf*TfR1 and SDDV. Our results demonstrate that *mf*TfR1 functions as a receptor for SDDV infection, specifically facilitating the binding of viral particles and clathrin-mediated endocytosis.

## RESULTS

### *mf*TfR1 is incorporated in purified SDDV viral particle

After treating the MFF-1-cultured SDDV suspension through gradient centrifugation, SDDV was well purified (Figure 1A) and *mf*TfR1 was identified in purified SDDV virion based on LC-MS/MS analysis (Table S1), with a total of five unique peptide matches (Figure 1B). As shown in Figure 1C, we have successfully generated a polyclonal antibody against the ectodomain of *mf*TfR1, which can specifically recognize *mf*TfR1 in its predicted location in SDS-PAGE gel. Furthermore, using virological methods combined with immunoblotting, *mf*TfR1 was found to localize to the nucleocapsid components of mature viral particles (Figure 1D). Among this component, the major capsid protein (MCP) of SDDV is predominantly localized in the pellet fraction, with minimal presence detected in the supernatant. As controls, SDDV-VP033 was previously identified as a minor capsid protein of SDDV (12), while SDDV-VP098 was determined to be a viral envelope protein (12), a homologous viral protein to ISKNV-VP56 (17).

**FIG. 1.**
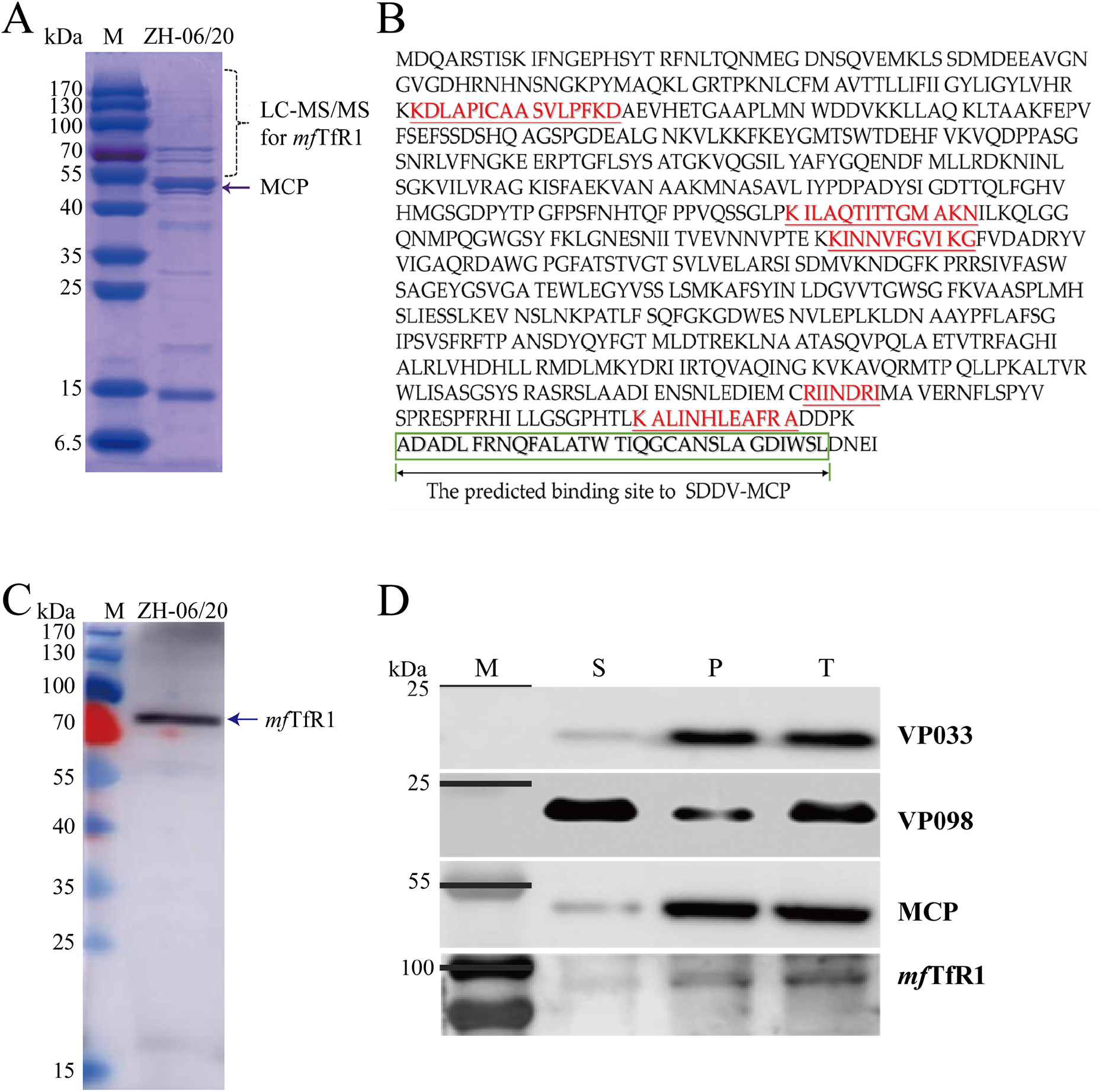
Identification of *mf*TfR1 as virion protein in purified SDDV. (A) Coomassie Brilliant Blue-stained gel electrophoresis image of purified SDDV ZH-06/20 virions; (B) Peptide matches of *mf*TfR1 in purified SDDV ZH-06/20 viral particles, with the matched peptides highlighted in red; (C) Specific recognition of *mf*TfR1 protein in ZH-06/20 virus particles by the *mf*TfR1 pAb; (D) Localization of *mf*TfR1 in purified SDDV virions. n = 3.

### *mf*TfR1 is involved in SDDV infection

To evaluate the potential role of *mf*TR1 in SDDV pathogenicity, we firstly analyzed its expression dynamics in mandarin fish. Real-time quantitative PCR (RT-qPCR) analysis demonstrated a marked reduction of *mf*TfR1 mRNA expression levels in the spleen and heart, where the highest levels of viral replication were detected. The downregulation of *mf*TfR1 mRNA expression became progressively more pronounced over time, reaching its lowest level at 14 dpi (Figure 2A). Complementary validation via immunofluorescence (IFA) and immunohistochemistry (IHC) in spleen tissues further corroborated the depletion of *mf*TfR1 protein in infected individuals (Figure 2B and C). Moreover, significant changes in *mf*TfR1 expression were observed across various tissues, providing preliminary evidence that *mf*TfR1 dynamically responds to SDDV infection. In addition, the relationship between *mf*TfR1 and SDDV infection in MFF-1 cells was investigated. As shown in Figure 2D, the relative expression of *mf*TfR1 continuously increased with infection time, which was also observed at the protein level (Figure 2E). Collectively, these findings implicate *mf*TR1 as a critical modulator of SDDV infection across *in vivo* and *in vitro* and that *mf*TfR1 is essential for SDDV infection. To establish the functional necessity of *mf*TfR1 in SDDV infection, we modulated its expression via overexpression and pharmacological inhibition. As shown in Figure 3A-C, overexpression of *mf*TfR1 in MFF-1 cells significantly increased *mf*TfR1 mRNA and protein levels, thereby enhancing SDDV infection. Further, fathead minnow cell (FHM), a low-permissive fish cell line to SDDV were transfected by pEGFP- *mf*TfR1, followed by infection with ZH-06/20 (MOI = 5) at 27 ∘. Strikingly, exogenous *mf*TfR1 expression rendered FHM cells permissive to SDDV, as evidenced by upregulated viral replication at transcriptional and translational levels (Figure 3D and E), which was validated by IFA analysis (Figure 3F). Ferristatin II, a compound known to induce *mf*TfR1 degradation (43), was employed to treat MFF-1 cells before SDDV infection. Cytotoxicity assays (CCK-8) confirmed the compound safety at 25–100 μM (Figure 3J). As shown in Figure 3K, Ferristatin II treatment significantly suppressed SDDV replication with a dose dependent effect. Using 50μM Ferristatin II to incubate cells, both mRNA and protein levels of *mf*TfR1 were significantly reduced (Figure 3L and M). When MFF-1 cells were pre-incubated with 50 μM of Ferristatin II and then were challenged with SDDV, WB analysis also revealed a reduction in SDDV protein levels (Figure 3N).

**FIG. 2.**
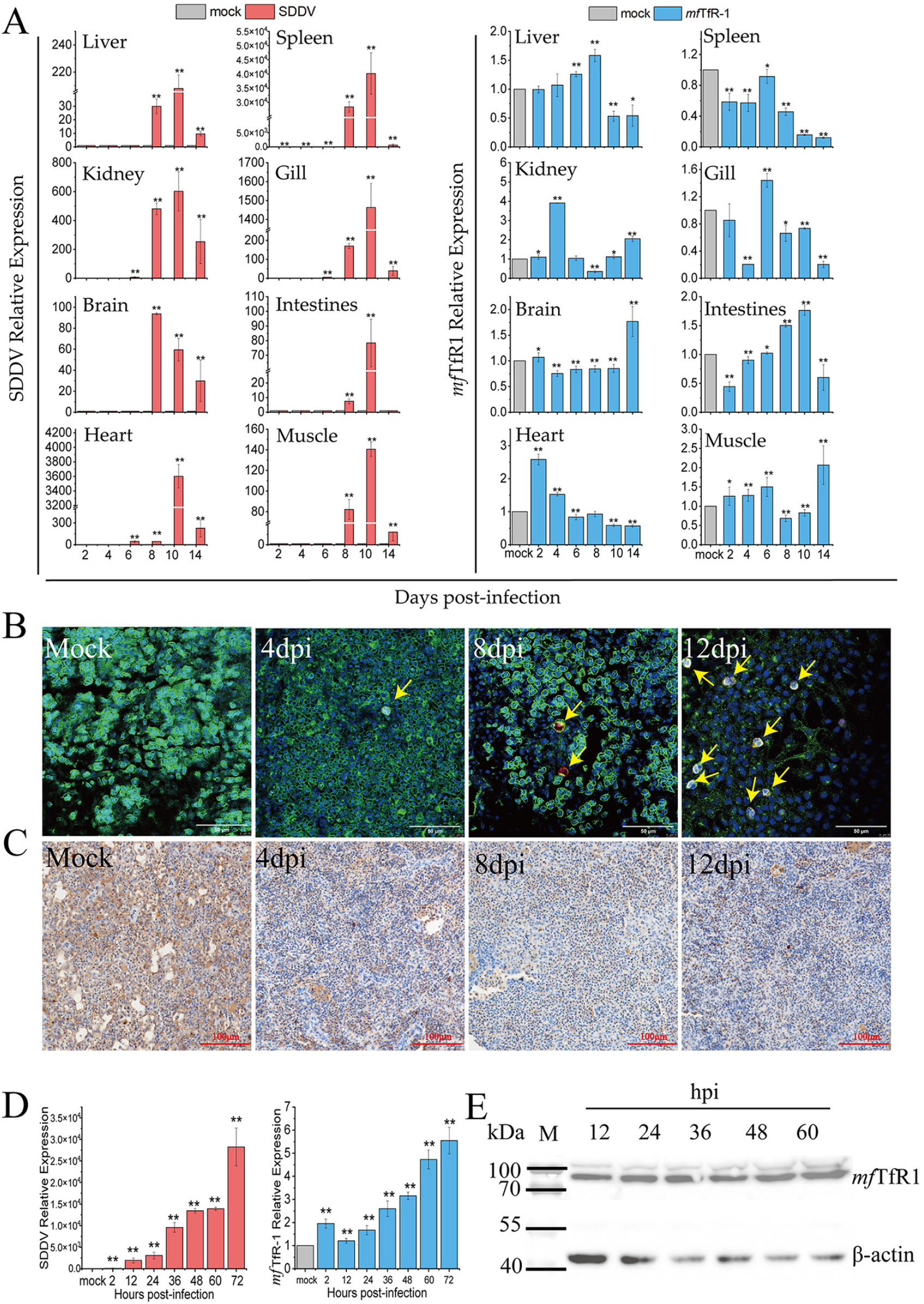
Expression of ***mf*TfR1** in response to SDDV infection *in vivo* and *in vitro*. (A) Changes in ***mf*TfR1** levels in various tissues (liver, spleen, kidney, gill, brain, intestine, heart and muscle) of mandarin fish infected with ZH-06/20. IFA (B) and IHC (C) detection of ***mf*TfR1** expression in the spleen. The yellow arrow points to the SDDV virus. (D) ***mf*TfR1** mRNA levels in MFF-1 cells after infection with ZH-06/20 was assessed by qPCR. (E) ***mf*TfR1** protein levels in MFF-1 cells after infection with ZH-06/20 was analyzed by Western blot. The data shown are means ± SD of three independent experiments or replicates. A two-tailed unpaired Student t test was used for the statistical analysis.n=3, \**p* < 0.05; ** *p* < 0.01.

**FIG. 3.**
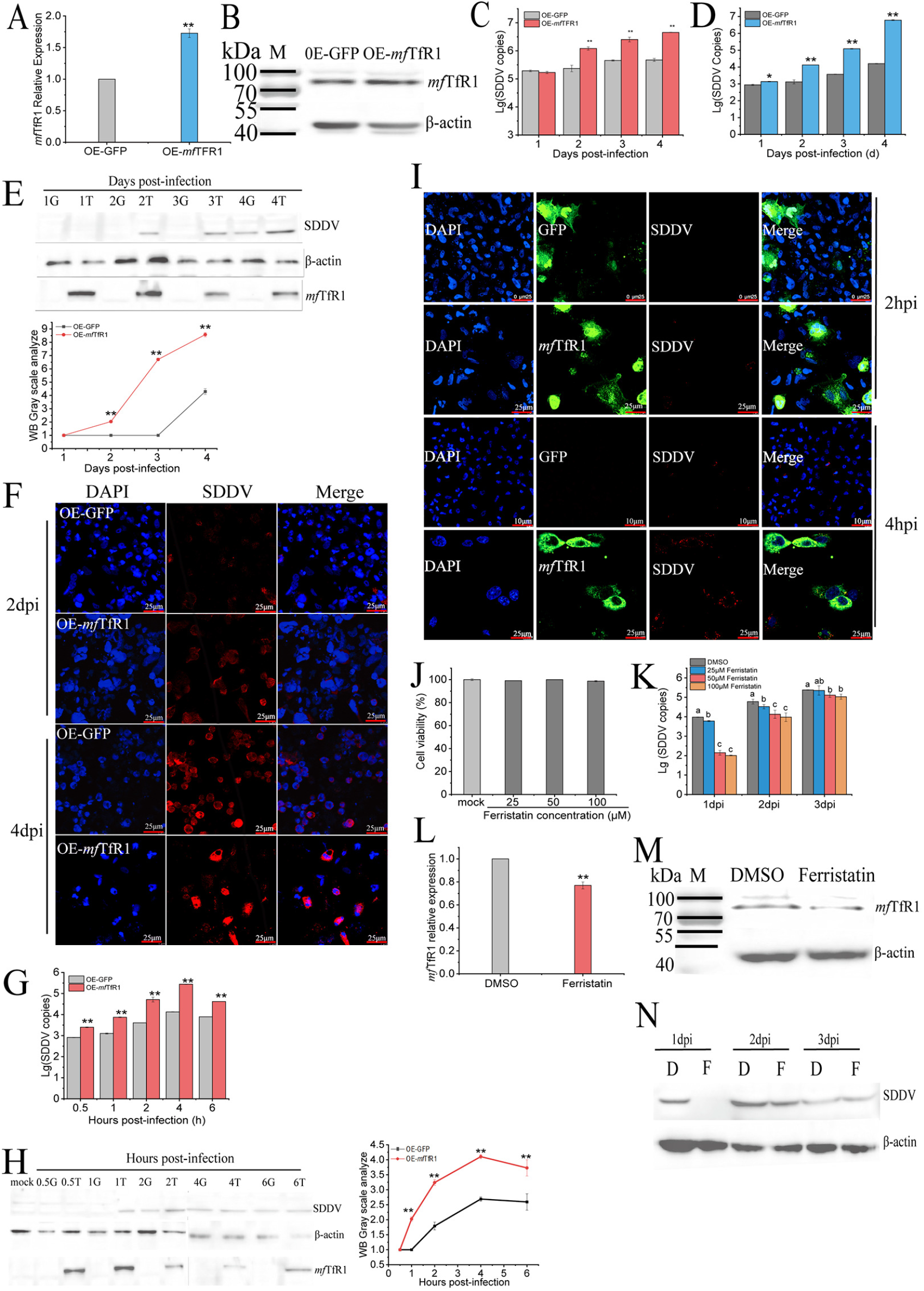
*mf*TfR1 is essential for SDDV infection. (A) mRNA levels in MFF-1 cells transfected with pEGFP- ***mf*TfR1** plasmid, compared to the control group transfected with pEGFP-N3. (B) Protein levels of ***mf*TfR1** in MFF-1 cells transfected with pEGFP- ***mf*TfR1**, compared to the control group. (C) After overexpression of ***mf*TfR1**, infection with ZH-06/20 resulted in a significant increase in SDDV copies compared to the pEGFP-N3 control group. (D) SDDV copies in FHM cells was significantly increased in the pEGFP- ***mf*TfR1 group** compared to the control group at 27°C from 1-4 dpi. (E) Western blot analysis of SDDV protein levels in FHM cells overexpressing ***mf*TfR1** at 27°C from 1-4 dpi, with gray scale analyze of Western blot bands (G represents OE-GFP and T represents OE- ***mf*TfR1**). (F) IFA analysis showing visual changes in SDDV infection at 2 dpi and 4 dpi at 27°C in ***mf*TfR1 group** compared to the OE-GFP group. (G) (D) SDDV copies in FHM cells was significantly increased in the pEGFP- ***mf*TfR1 group** compared to the control group at 4°C from 0.5-6 dpi. (H) Western blot analysis of SDDV protein levels in FHM cells overexpressing ***mf*TfR1** at 4°C (0.5-6 hpi), with gray scale analyze of Western blot bands. (I) IFA analysis showing visual changes in SDDV infection at 2 hpi and 4 hpi at 4°C in ***mf*TfR1** overexpressing cells compared to the OE-GFP group. (J) CCK-8 assay showing no effect of ferristatin II (25-100 μM) on cell viability. (K) SDDV copies in cells pretreated with different concentrations of ferristatin II before infection (1-3 dpi). (L, M) mRNA (qPCR) and protein (Western blot) analysis showing significant reduction in ***mf*TfR1** levels following 50 μM ferristatin II treatment. (N) Western blot analysis of SDDV protein levels in cells pretreated with 50 μM ferristatin II. D indicates the DMSO group, and F indicates the ferristatin II group. The data shown are means ±SD of three independent experiments or replicates. A two-tailed unpaired Student t test was used for the statistical analysis. n=3, \**p* < 0.05; \*\**p* < 0.01. Different letters (a, b, c, d) indicate significant differences between groups.

Further, FHM cells overexpressed with *mf*TfR1 were placed on ice and infected with SDDV (MOI = 25). At this temperature, the virus could bind to cells without entering the cells. As shown in Figure 3G, overexpression of *mf*TfR1 (OE- *mf*TfR1) in FHM cell significantly facilitated SDDV binding. This trend was further confirmed by Western blot analysis (Figure 3H). According to IFA, in the OE- *mf*TfR1 group, red fluorescence representing SDDV was first observed at 2 hpi and intensified by 4 hpi, while only weak red fluorescence was detected in the control group at 4 hpi (Figure 3I). These results indicated that *mf*TfR1 plays a critical role in the process of SDDV infection, especially entering the cells.

### *mf*TfR1 co-localizes with SDDV via interacting with SDDV MCP

MCP is the most abundant protein in iridovirus, accounting for approximately 40-50% of the viral structural proteins (17, 18). Therefore, we intend to explore whether *mf*TfR1 interacts with SDDV MCP. First, IFA of spleen tissues from SDDV-infected mandarin fish revealed that in uninfected cells, *mf*TfR1 was localized to the cell membrane. However, following SDDV infection, the nuclei of SDDV-infected cells appeared enlarged, and *mf*TfR1 was induced to translocate into the cytoplasm and co-localized with SDDV MCP (Figure 4A). Similarly, in SDDV-infected MFF-1 cells, MCP also co-localized with *mf*TfR1 (Figure 4A). And a high degree of colocalization between *mf*TfR1 and MCP was observed in pEGFP-*mf*TfR1 and pmCherry-MCP plasmids co-transfected FHM cells (Figure 4A). To validate physical interaction, co-immunoprecipitation (Co-IP) assays were conducted, demonstrating the reciprocal binding between *mf*TfR1 and MCP (Figure 4B and C). These findings collectively establish *mf*TfR1 as a direct interactor of SDDV MCP during SDDV infection.

**FIG. 4.**
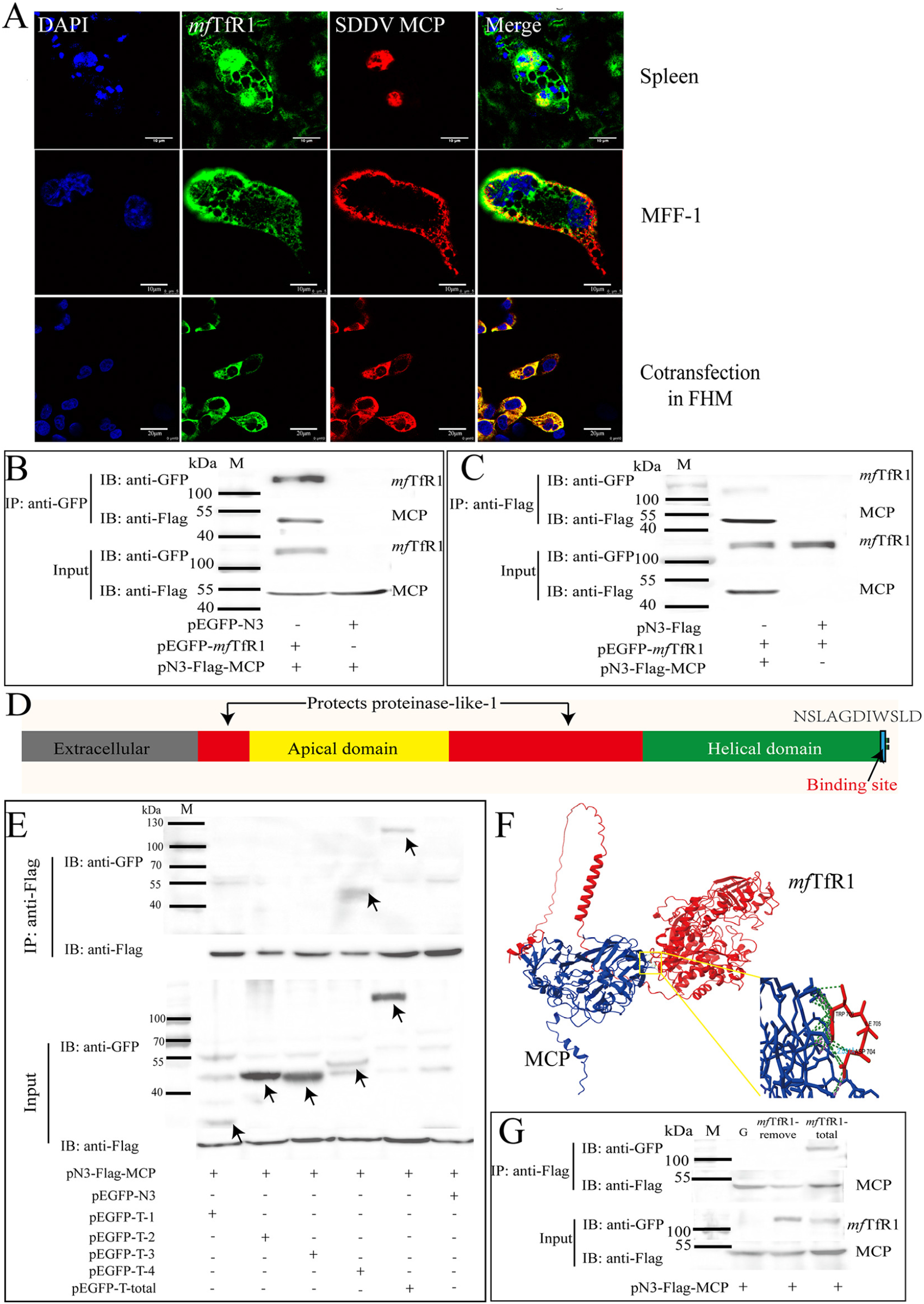
*mf*TfR1 directly interacts with SDDV MCP. (A) Localization of ***mf*TfR1** and MCP in mandarin fish spleen tissue, MFF-1 cells after SDDV infection, and FHM cells co-transfected with ***mf*TfR1** and MCP. (B, C) FHM cells were co-transfected with pEGFP-***mf*TfR1** and pFlagN3-SDDV MCP, followed by Co-ip using anti-Flag agarose beads (B) or anti-GFP agarose beads (C). (D) Mapping of ***mf*TfR1** protein domains and the binding region with MCP. (E) Co-transfection of ***mf*TfR1**-GFP and various ***mf*TfR1** truncation mutants (T-1-GFP, T-2-GFP, T-3-GFP, T-4-GFP) with SDDV MCP-Flag, followed by Co-ip using anti-GFP agarose beads. (F) Binding site prediction of ***mf*TfR1** and MCP proteins by Chimera X software. (G) Co-transfection of ***mf*TfR1**-deletion mutant and SDDV MCP-Flag, followed by Co-ip using anti-Flag agarose beads. The data shown are means ±SD of three independent experiments or replicates. A two-tailed unpaired Student t test was used for the statistical analysis. n=3, \**p* < 0.05; \*\**p* < 0.01.

By comparing with structural domains of human TfR1 (*h*TfR1) and mouse TfR1 (*m*TfR1), we can divide the extracellular domain of TfR1 into the protease-like domain, apical domain, and helical domain, respectively (Figure 4D). To determine the specific domain which may interact with MCP, EGFP-tagged distinct domains of *mf*TfR1 were constructed, including Protease-like domain-1 (T-1-GFP), Apical domain (T-2-GFP), Protease-like domain-2 (T-3-GFP) and Helical domain (T-4-GFP). FHM cells were co-transfected with pFlag-N3-SDDV MCP and these different recombinant *mf*TfR1 domain plasmids, respectively, followed by Co-IP assays. As a result, pEGFP-*mf*TfR1-Helical domain was associated with pFlag-N3-SDDV MCP, whereas no interaction was observed between pEGFP-*mf*TfR1-Protease-like domain-1, pEGFP- *mf*TfR1-Apical domain, pEGFP-*mf*TfR1-Protease-like domain-2 and pFlag-N3-SDDV MCP (Figure 4E), revealing that the helical domain of *mf*TfR1 is responsible for binding to MCP. To further identify the precise binding sites between *mf*TfR1 and MCP, Chimera X software was used to predict the interaction interface of the two proteins. As shown in Figure 4D and F, the binding sites were identified as the tryptophan (TRP) and aspartic acid (ASP) residues at the C-terminus of *mf*TfR1. A mutant plasmid, lacking the C-terminal site including the predicted binding site (pEGFP- *mf*TfR1-remove), was constructed and co-transfected with pFlag-SDDV MCP into FHM cells. Co-IP results showed that pEGFP-*mf*TfR1-remove plasmid completely abolished the interaction between *mf*TfR1 and MCP (Figure 4G). Collectively, these findings highlight the critical role of the helical domain of *mf*TfR1 and its specific residues in mediating the interaction with MCP.

### *mf*TfR1 is an entry receptor for SDDV

*mf*TfR1 functions as a critical entry receptor facilitating SDDV infection. To validate this role, a neutralizing antibody targeting the extracellular domain of *mf*TfR1 and a polypeptide (TM-B) containing the binding site (sequence: ADL FRN QFA LAT WTI QGC ANS LAG DIW SL) with SDDV-MCP in *mf*TfR1 were used to evaluate the role of *mf*TfR1 in the process of SDDV infection. Cell viability was unaffected by antibody and TM-B treatment (Figure 5). As shown in figure 5A, antibody treatment significantly inhibited CPE induced by SDDV infection in a dose-dependent manner. And blocking *mf*TfR1 significantly reduced expression levels of *mcp* (Figure 5B). IFA corroborated these findings, showing diminished viral signal intensity in antibody-treated groups (Figure 5C). Additionally, TM-B polypeptide significantly decreased SDDV infection (Figure 5D) and CPE was obviously weakened by TM-B treatment (Figure 5E). These results were further confirmed by IFA analysis (Figure 5F). And impeded viral entry into cytoplasmic and nuclear compartments, as visualized by fluorescence tracking (Figure 5F). When mandarin fish was challenged with TM-B polypeptide and SDDV mixtures, a relatively lower mortality rate and delayed death time compared to the control group treated with PetB2M protein and SDDV mixtures was observed (Figure 5G). Consistently, qRT-PCR analysis also confirmed a marked reduction in the relative expression of SDDV *mcp* in TM-B treatment group (Figure 5H). In summary, the results confirm that *mf*TfR1 is a key receptor of SDDV infection.

**FIG. 5.**
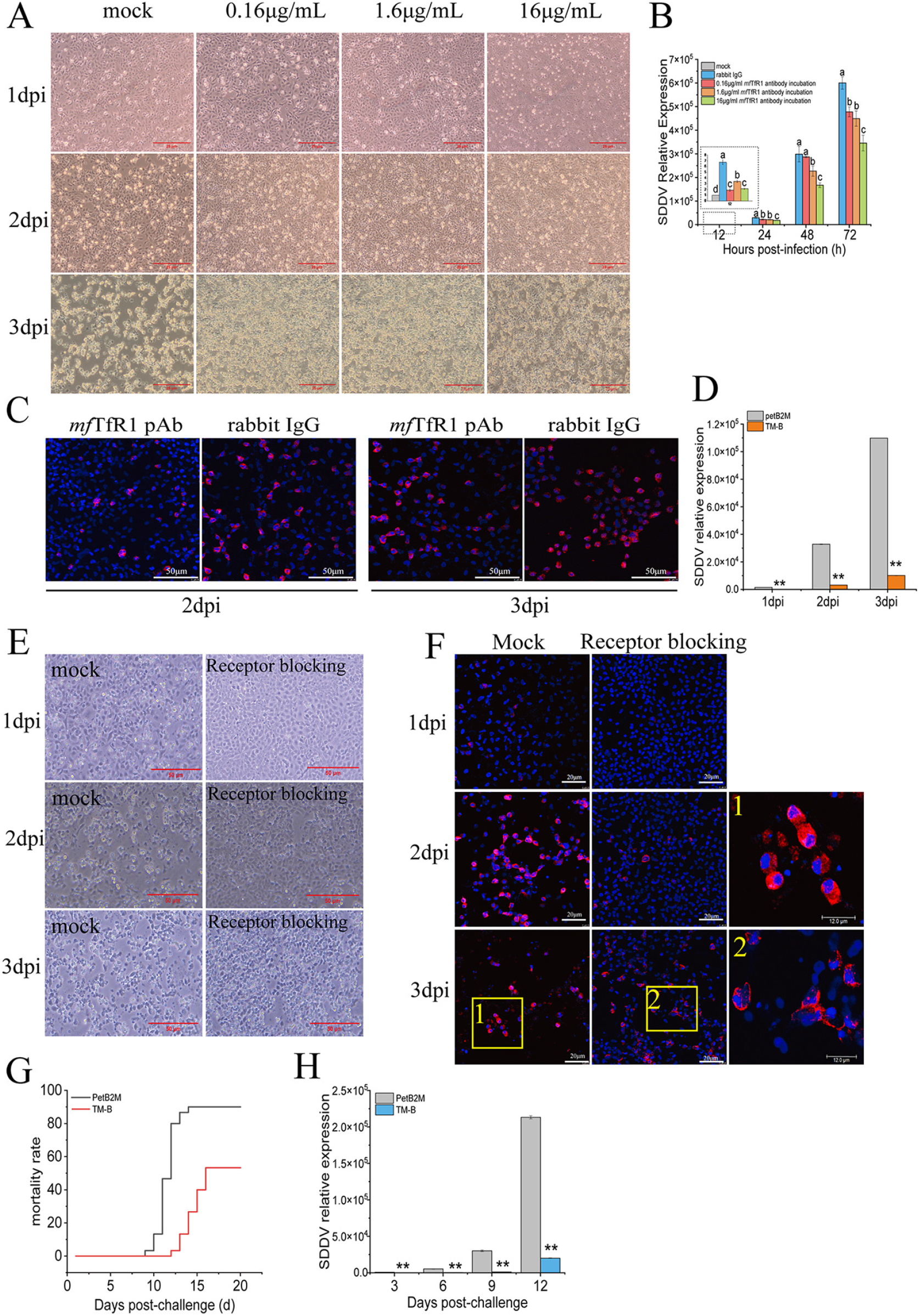
*mf*TfR1 is an entry receptor for SDDV. (A) CPE observed in cells treated with 0.16, 1.6, and 16 μg/ml ***mf*TfR1** pAb followed by SDDV infection. (B) qPCR results showing the reduction in SDDV mRNA levels in cells pretreated with ***mf*TfR1** pAb. Significant decrease in ***mf*TfR1** mRNA (C) IFA showing changes in SDDV protein levels after ***mf*TfR1** pAb treatment compared to the control group. ZH-06/20 was pooled with TM-B (50μg/ml), The petB2M protein was used as the negative control. SDDV was then detected by qPCR (D), CPE observation (E), and IFA (F). (G) Mortality rate of mandarin fish injected with a mixture of TM-B and ZH-06/20. The petB2M protein was used as a negative control. (H) Relative SDDV expression in the spleen tissues of infected mandarin fish, measured by qPCR. The data shown are means ±SD of three independent experiments or replicates. A two-tailed unpaired Student t test was used for the statistical analysis. n=3, \**p* < 0.05; \*\**p* < 0.01.

### *mf*TfR1 is required for the endocytosis of SDDV

Following receptor engagement by viral structural proteins, host cells typically activate endocytic mechanisms to facilitate viral entry. To determine whether SDDV enters cells via *mf*TfR1-mediated endocytosis, we firstly examined the role of CME in SDDV entry. Membrane cholesterol, a key regulator of clathrin-coated pit assembly (44, 45), was pharmacologically modulated using, Methyl-β-cyclodextrin (MβCD, a cholesterol-depleting agent) and Nystatin (a cholesterol-sequestering compound) (46). MFF-1 cells were pretreated with graded concentrations of MβCD or Nystatin, exposed to SDDV at 4°C for 1 h to permit viral attachment, and then shifted to 27 °C for 2.5 h to initiate internalization. Trypsin treatment was applied to eliminate surface-bound SDDV. qPCR results showed MβCD had no effect on SDDV binding to the cell membrane but significantly inhibited viral internalization (Figure 6B and C). Conversely, Nystatin promoted SDDV internalization without affecting binding (Figure 6B and C). These findings indicate that SDDV enters cells through endocytosis.

**FIG. 6.**
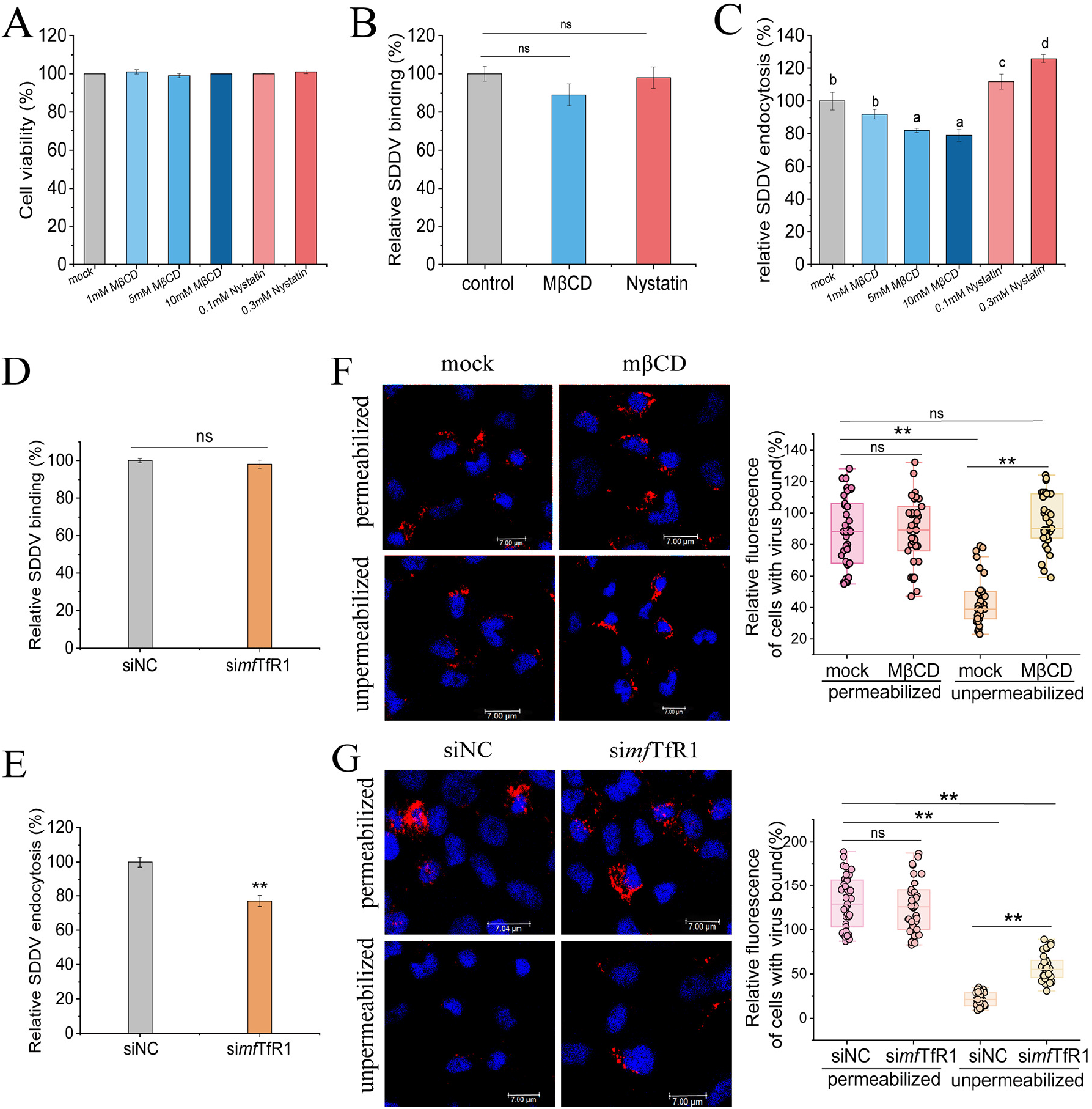
*mf*TfR1 is required for the endocytosis of SDDV. (A) CCK-8 assay showing no effect of MβCD or Nystatin on cell viability. SDDV binding (B) and entry (C) assays were performed with MβCD or Nystatin-treated cells. Viral binding or entry was quantified by normalization to the respective control cells. RABV binding (D) and entry (E) assays were performed with ***mf*TfR1**-silenced MFF-1 cells. Viral binding or entry was quantified by normalization to the control cells. (F) MβCD-treated cells were processed as described for panel E, except that they were not treated with trypsin. SDDV was stained with the antibody against SDDV MCP (red). Cell nuclei were stained with Hoechst 33342. Representative images are shown. The fluorescence signal of SDDV particles was quantified by using Image J software. The relative fluorescence of cell-bound SDDV under permeabilized or unpermeabilized conditions was quantified by normalization to the control cells under permeabilized conditions. The circles represent individual data points. At least 35 cells per sample were quantified. (G) ***mf*TfR1**-silenced cells were processed as described for panel F. In panels A to E, the data shown are means ± SD of three independent experiments or replicates. In panels F and G, the data represent the sum of three independent experiments. A two-tailed unpaired Student t test was used for the statistical analysis. ns, not significant; n=3, \**p* < 0.05; ** *p* < 0.01. Different letters (a, b, c, d) indicate significant differences between groups.

To investigate the functional role of *mf*TfR1 in SDDV entry, we used RNA interference (RNAi) to examine its effect on viral binding and internalization. Silencing *mf*TfR1 (si *mf*TfR1) in MFF-1 cells did not affect SDDV binding (Figure 6D) but severely impaired intracellular uptake (Figure 6E), highlighting its essential role for the endocytosis of SDDV. To further dissect this process, a microscopy-based assay was developed: control and si*mf*TfR1 group were incubated with SDDV MCP mAb under permeabilized or non-permeabilized conditions, followed by IFA. Non-permeabilized cells reflect surface-bound virions (internalized particles inaccessible to antibodies), whereas permeabilized cells reveal total viral load. Cholesterol depletion via MβCD served as a positive control for impaired endocytosis. In si*mf*TfR1 group, non-permeabilized fluorescence intensity exceeded controls (Figure 6F), indicating defective internalization, while permeabilized measurements showed comparable viral accumulation between groups (Figure 6G). These data conclusively position *mf*TfR1 as a critical mediator of SDDV endocytosis, independent of initial viral binding.

### Src mediates *mf*TfR1 internalization during SDDV infection

TfR1 internalization is regulated by Src kinase (47). During the first 0.5 h of SDDV infection, the total levels of *mf*TfR1 and Src in MFF-1 cells remained unchanged (Figure 7A). To investigate the mechanism of *mf*TfR1-mediated SDDV entry via CME, the membrane domain of *mf*TfR1 was extracted from SDDV-infected cells at 0.5 hpi. A decrease in membrane-associated *mf*TfR1 was observed (Figure 7B), indicative of activated receptor endocytosis. To characterize the molecular interplay between *mf*TfR1 and Src kinase during SDDV infection, lysates from infected MFF-1 cells (0.5–4 hpi) were subjected to immunoprecipitation (IP) with antibodies targeting *mf*TfR1 or Src, followed by subsequent immunoblot analysis. IP assays revealed a stable association between *mf*TfR1 and Src, with SDDV infection markedly enhancing this interaction (Figure 7C). revealed that Src binds to *mf*TfR1 and that SDDV infection enhanced the interaction between *mf*TfR1 and Src. Moreover, *mf*TfR1 tyrosine phosphorylation was detected in SDDV-infected cells. Immunoblotting results (Fig. 7C and D) demonstrated that SDDV infection induced an increase in tyrosine phosphorylation (p-Tyr) of *mf*TfR1. These observations suggest that Src and *mf*TfR1 form a stable complex following infection. To determine whether Src is critical for *mf*TfR1-mediated SDDV entry, MFF-1 cells were pretreated with the Src inhibitor PP2 prior to infection. RT-qPCR analysis revealed that Src inhibition significantly impaired the ability of *mf*TfR1 to mediate SDDV infection (Fig. 7E).

**FIG. 7.**
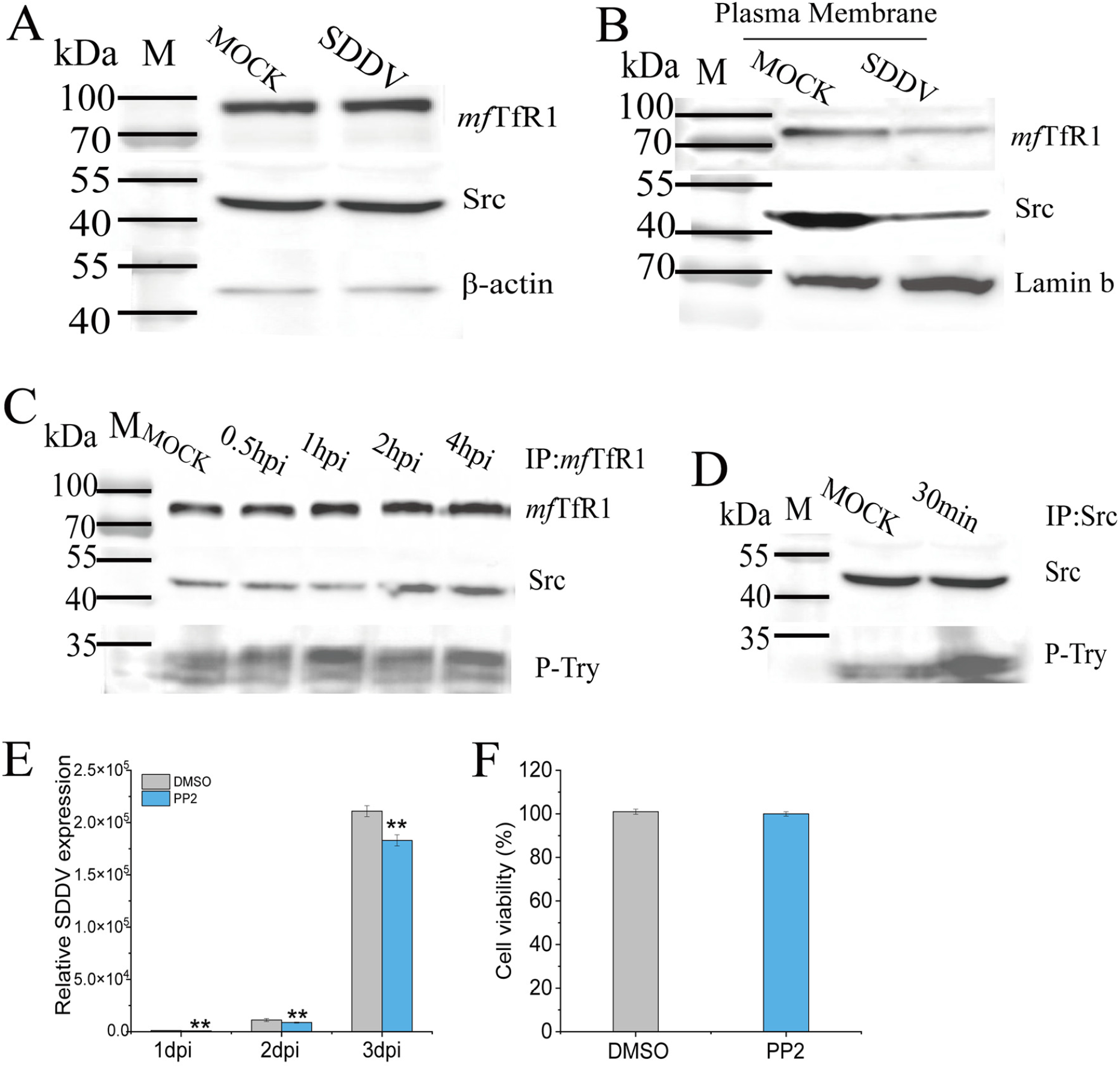
*mf*TfR1 internalization is associated with Src during SDDV infection. (A) Western blots of ***mf*TfR1** and Src levels in SDDV infected MFF-1 cells. (B) Western blots of membrane ***mf*TfR1** and Src in infected cells. (C) Western blot of the immunoprecipitation of ***mf*TfR1**, p-Tyr, and Src from SDDV infected cells. (D) Western blot of the immunoprecipitation of p-Tyr and Src from SDDV infected cells. (E) mRNA level of internalized SDDV in MFF-1 cells pretreated with the Src inhibitor PP2 then subsequently infected. (F) CCK-8 assay showing no effect of PP2 on cell viability. E, the data shown are means ± SD of three independent experiments or replicates. A two-tailed unpaired Student t test was used for the statistical analysis. n=3, * *p* < 0.05; ** *p* < 0.01.

## Discussion

Compared to the extensively studied ISKNV, SDDV exhibits considerable differences in its genome organization, protein composition, serotype, and pathogenicity (2, 3, 13). Despite its increasing relevance, the pathogenic mechanisms underlying SDDV infection remain largely unexplored. Our previous studies revealed that SDDV infection not only enhances systemic iron storage but also induces ferroptosis both *in vivo* and *in vitro* (22). *mf*TfR1, a pivotal regulator of cellular iron homeostasis, was identified as a key mediator of SDDV-induced ferroptosis (22). In this study, we further detected abundant *mf*TfR1 protein within purified SDDV virions, indicating a possible interaction between viral structural protein and *mf*TfR1. Tissue-specific mRNA analysis demonstrated a ubiquitous expression of *mf*TfR1 across various tissues in SDDV-infected mandarin fish, with the most pronounced downregulation observation in spleen—the most important target organ of SDDV infection (2, 21). These findings were further validated by IFA and IHC, reinforcing the hypothesis that *mf*TfR1 plays a critical role in SDDV pathogenesis. Intriguingly, *in vitro* experiments using SDDV-infected MFF-1 cells revealed a stark contrast, with *mf*TfR1 expression significantly upregulated. The expression of *mf*TfR1 is governed by a tightly regulated negative feedback loop within the iron homeostasis pathway. Elevated intracellular iron levels typically lead to the downregulation of *mf*TfR1, reducing cellular iron uptake (48). In spleen, which serves as a major iron reservoir, SDDV infection may induce iron overload, triggering the downregulation of *mf*TfR1 as a defensive mechanism to mitigate excessive iron uptake. In contrast, the *in vitro* studies suggest that SDDV may exploit *mf*TfR1 as a receptor to facilitate cellular entry. By upregulating *mf*TfR1 expression, the virus appears to enhance its infectivity and invasiveness. The observed discrepancies between *in vivo* and *in vitro* findings underscore the intricate complexity of host-virus interactions, may suggesting a dual role for *mf*TfR1 in SDDV infection: functioning both as a suppressor of ferroptosis and as a critical mediator facilitating viral entry.

The hypothesis derived from the above findings is supported by *in vitro* results, including *mf*TfR1 overexpression, and ferristatin II treatment. In MFF-1 cells, overexpression of *mf*TfR1 can promote viral replication. FHM cells, which are typically less sensitive to SDDV, exhibited significantly enhanced SDDV replication following *mf*TfR1 overexpression. Moreover, SDDV infection in FHM cells was analyzed in two stages: viral binding and entry. By comparing the infection trend slopes between the OE-GFP and OE- *mf*TfR1 groups, it was observed that the slopes became parallel during viral binding (2–4 hpi) and entry (3–4 dpi). This indicates that *mf*TfR1 primarily affects the early stages of viral binding and entry, consistent with Zhang et al.’s findings on Nectin-1-mediated RGNNV entry (49).

Virus entry is the first crucial step in infection, and determining the associated binding proteins and their specific cellular entry sites plays a pivotal role, both in elucidating the underlying mechanisms and in guiding potential therapeutic interventions. MCP of SDDV forms the structural backbone of the viral capsid, serving as the primary protective shell for its genomic material. WB analysis of purified SDDV virions revealed a predominant localization of both MCP and *mf*TfR1 in the cellular pellet fraction (Fig. 1D). IFA performed on spleen tissues demonstrated a striking colocalization of SDDV MCP and *mf*TfR1 (Fig. 4A). Notably, their localization shifted progressively from the cell membrane to the cytoplasm and nucleus as the infection advanced. These findings suggest the dynamic interactions between MCP and *mf*TfR1 during SDDV infection. In MFF-1 cells infected with SDDV and FHM cells co-transfected with pEGFP-*mf*TfR1 and pmCherry-MCP, *mf*TfR1 and MCP exhibited clear colocalization, revealing the important role of *mf*TfR1 in the SDDV infection process and its interaction with SDDV MCP. Further investigation through Co-IP assays confirmed the specificity of the MCP- *mf*TfR1 interaction.

The structure of hTfR1 and mTfR1 has been elucidated through X-ray crystallography and cryo-electron microscopy (cryo-EM), including both the free-state TfR1 structure and its complex with transferrin (50). TfR1 forms a butterfly-shaped extracellular domain dimer, which consists of a stalk domain approximately 30 amino acids long, a transmembrane sequence, and an N-terminal cytoplasmic tail. The cytoplasmic tail interacts with Adaptor Protein 2 (AP2) and mediates CME (50). Each monomer of the extracellular domain of TfR1 contains three major structural domains: the protease-like domain near the membrane, the helical domain that forms the dimer interface, and the apical domain at the distal end. To determine the binding domains and sites of *mf*TfR1 and MCP, we firstly predicted the structural domains of *mf*TfR1 using <u>AlphaFold3,</u> SnapGene software, and Chimera X software. Subsequently, we successfully obtained the predicted binding sites by analyzing the binding conformation of *mf*TfR1 and MCP in <u>AlphaFold3</u> and visualizing them in Chimera X software. Subsequent Co-IP experiments precisely validated the interaction regions between these two proteins, identifying the helical domain of *mf*TfR1 as the specific docking site for MCP.

TfR1 is ubiquitously expressed across tissues and is indispensable for cellular function, with genetic ablation being universally lethal (51, 52). This biological constraint precludes the development of TfR1-knockout mandarin fish to study its role in SDDV pathogenesis. However, the identification of the MCP- *mf*TfR1 binding interface opens avenues for the creation of domain-deleted mandarin fish models, which could provide a powerful tool to dissect the role of *mf*TfR1 *in vivo*. Despite the absence of reported gene knockout studies in mandarin fish, foundational research has established a robust framework. Recent advances, including chromosome-level genome sequencing of mandarin fish, have provided critical insights into key traits such as aggressive feeding, growth, pyloric caecum development, and salinity adaptation (53). These genomic resources lay the groundwork for future gene-editing initiatives to explore the molecular basis of SDDV-host interactions. In the present study, antibody-blocking and receptor-blocking experiments further substantiated the pivotal role of *mf*TfR1 in SDDV infection. Incubation with *mf*TfR1 pAb in MFF-1 cells effectively reduced SDDV replication. In addition, High-purity (>95%) TM-B were synthesized and used to pre-incubate SDDV virions and MFF-1 cells, respectively. Both *in vitro* and *in vivo* results revealed a significant reduction in SDDV infectivity. Notably, mandarin fish challenged with SDDV pre-incubated with TM-B exhibited a remarkable 40% reduction in mortality. These findings strongly implicate *mf*TfR1 as a receptor mediating SDDV infection and highlight its potential as a therapeutic target for controlling the disease.

Iron-bound transferrin (Tf) is internalized by TfR1 through clathrin-mediated endocytosis within clathrin-coated pits (54). Treatment of cells with MβCD and Nystatin followed by SDDV challenge demonstrated that SDDV utilizes an endocytic pathway for cell entry. Furthermore, silencing TfR1 confirmed its essential role in the endocytosis of SDDV. Src tyrosine kinase, a critical regulator of cell signaling pathways (55), has been shown to mediate the endocytosis of TfR1 through its activation (56). In agreement with findings by Jian et al. (47), our results demonstrated that *mf*TfR1 interacts with Src and that SDDV infection enhances Src activation, leading to Tyr^20^ phosphorylation of *mf*TfR1. Notably, the Src inhibitor PP2 significantly suppressed SDDV infection. These findings suggest that Src facilitates *mf*TfR1 phosphorylation, driving its endocytosis and thereby promoting SDDV cellular entry.

In conclusion, our study establishes *mf*TfR1 as a critical receptor for SDDV entry (Fig. 8), with MCP-*mf*TfR1 interactions driving viral binding, endocytosis, and replication. Structural and functional analyses, along with receptor-blocking and peptide inhibition assays, confirm that disrupting this interaction significantly reduces SDDV infectivity and host mortality (Fig. 8). These findings enhance our understanding of SDDV pathogenesis and position *mf*TfR1 as a promising therapeutic target. Future efforts should focus on developing peptide-based inhibitors or small molecules to block *mf*TfR1-mediated viral entry and leveraging genetic tools to clarify its *in vivo* role. Such advances will facilitate effective strategies to control SDDV in aquaculture.

**FIG. 8.**
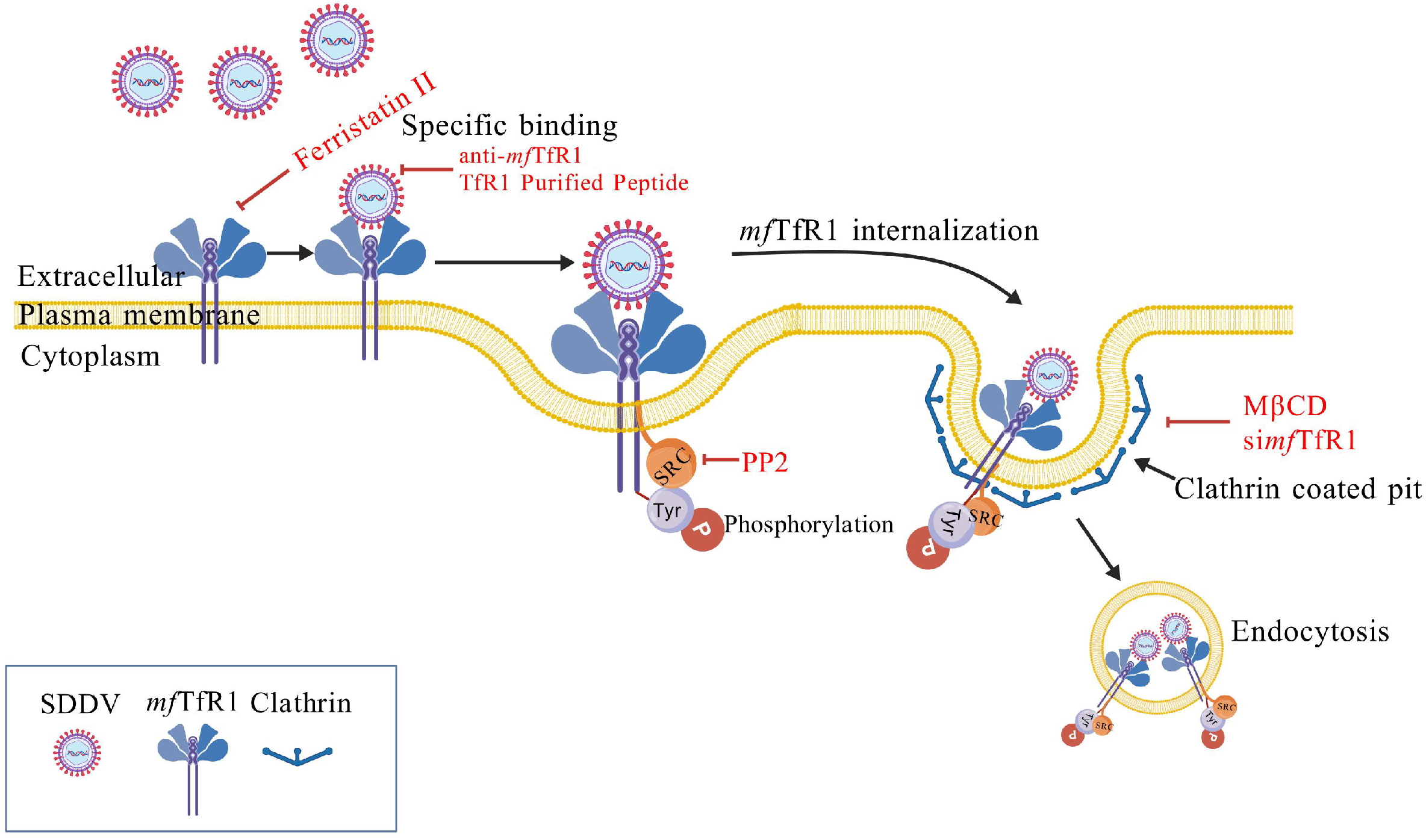
The schematic of *mf*TfR1 is an entry factor for SDDV mediating viral entry via clathrin-mediated endocytosis. During infection, SDDV specifically interacts with *mf*TfR1, leading to the activation of Src kinase, which subsequently induces tyrosine phosphorylation of *mf*TfR1 and promotes its internalization into host cells via CME. The image is created with BioGDP.com (https://biogdp.com, BioGDP Agreement Number: GDP2025IWMEE4).

## MATERIALS AND METHODS

### Cell lines, viruses and antibodies

Mandarin fish fry (MFF-1) cell line (57) and FHM cells (58) were established and maintained in our laboratory, as served as primary models for viral isolation and infection assays. MFF-1 cells were propagated in Dulbecco’s Modified Eagle’s Medium (DMEM), while FHM cells were grown in Medium 199 (M199), both media containing 10% fetal bovine serum (FBS), 100 IU/mL penicillin, 100 μg/mL streptomycin, and 0.25 μg/mL amphotericin B (Life Sciences, USA) under 27 °C incubation. The SDDV strain ZH-06/20, was originally isolated and characterized in our group, growing in MFF-1 cells and being preserved at −80 °C until use (2).

Rabbit polyclonal anti- ***mf*TfR1** (***mf*TfR1** pAb) was prepared for this study, briefly, genomic DNA from mandarin fish tissue was used as the template. The PCR product was purified and cloned into the pMal-c2X vector to construct the pMal-c2X-MCP plasmid. Sequencing confirmed the recombinant plasmid, which was expressed in Escherichia coli BL21. After IPTG induction (1 mM at 37 °C for 6 h), the MBP- ***mf*TfR1** fusion protein was expressed and analyzed by SDS-PAGE. Bacterial cells were collected, lysed, and centrifuged. Proteins in the supernatant and pellet were analyzed by SDS-PAGE, and concentrations were determined using a BCA Protein Assay Kit (TaKaRa, Japan). The MBP- ***mf*TfR1** fusion protein was excised from Coomassie-stained gels, ground in PBS, and emulsified with Freund’s adjuvants for immunization. New Zealand rabbits received four subcutaneous injections at 2-week intervals, and serum was collected two weeks after the final immunization and stored at −80 °C. Mouse monoclonal anti-SDDV MCP protein (MCP mAb) and rabbit polyclonal anti-SDDV VP033 and VP098 protein was prepared and characterized in our laboratory (12). Anti-β-actin monoclonal antibody (Merck millipore, Germany). Mouse monoclonal antibodies to flag and GFP (Mabstar, China). Anti-Phosphotyrosine polyclonal antibody (Abcam, Britain). Anti-Src polyclonal antibody (Src pAb) (Proteintech, USA). HRP-conjugated goat anti-rabbit and anti-mouse IgG (Promega, USA). Alexa Fluor 555-conjugated donkey anti-mouse IgG (Invitrogen, USA). Alexa Fluor 488-conjugated donkey anti-rabbit IgG (Invitrogen, USA).

### Plasmids construction

Plasmid constructs were engineered through established molecular cloning protocols. Target sequences were amplified by PCR, followed by restriction enzyme digestion and homology-directed assembly. DNA fragments were subsequently inserted into appropriate vectors using the 2× MultiF Seamless Assembly Kit (ABclonal, China). ***mf*TfR1** full-length (fl) (Accession No.: MK605398.1), apical domain (ad), protease-like domain (pld), helical domain (hd), and the residual sequence excluding the ligand-binding site (remove) were cloned into pEGFP-N3 vectors. Additionally, the SDDV MCP cDNA (Accession No.: MH152405.1) was cloned into pFlag-N3 vectors, as described in this study. All constructs were validated by sequencing analysis. Primers RT-PCR were designed using NCBI Primer-BLAST (**Table S2**).

### Virus purification

The culture method of SDDV ZH-06/20 strain is as described above. Virus-infected cells were harvested at 72 hpi when significant CPE were observed. The cell culture supernatant was clarified by centrifugation at 4,000 × g for 15 min at 4 °C to remove cell debris. The supernatant was then ultracentrifuged at 100,000 × g for 2 h at 4 °C using a 30% sucrose cushion. The viral pellet was resuspended in PBS and further purified by a discontinuous sucrose gradient (40%, 50%, and 60%) via ultracentrifugation at 100,000 × g for 2 h at 4 °C. Purified virus particles were collected from the 50–60% interface, washed with PBS, and stored at −80 °C until analysis.

### LC-MS/MS

Protein identification was performed following standard protocols as our previous report (17, 18). LC-MS/MS analysis was conducted by Shenzhen Gene Genius Biotechnology Co., Ltd.

### Western blotting

Protein samples from MFF-1 cells were extracted using RIPA lysis buffer (Thermo Fisher Scientific, USA). Equal protein quantities were separated on 10% SDS-PAGE gels and transferred to PVDF membranes (Merck Millipore, Germany). Membranes were probed with primary antibodies: *mf*TfR1 pAb (1:1,000), MCP mAb (1:1000) and Src pAb (1:500). Loading controls was β-actin (mouse monoclonal, 1:1,000). HRP-conjugated goat anti-rabbit/mouse IgG served as the secondary antibody. Using High-sig ECL substrate to detect protein bands (Tanon, China).

### RT-qPCR

primers are listed in Table S3. The assay was performed according to established protocols (22).

### IHC and IFA

Antibody-based IHC and IFA were performed to detect SDDV and ***mf*TfR1**. Mouse anti-MCP mAb and rabbit anti-***mf*TfR1** pAb (1:500 dilution) were used as primary antibodies. For IHC, horseradish peroxidase (HRP)-labeled goat anti-mouse IgG (against SDDV MCP) or HRP-labeled goat anti-rabbit IgG (against ***mf*TfR1**) served as secondary antibodies. Tissue sections were developed using 3,3′-diaminobenzidine (DAB) solution for visualiz ation. For IFA, Alexa Fluor 555-conjugated donkey anti-mouse IgG (against SDDV MCP) and Alexa Fluor 488-conjugated donkey anti-rabbit IgG (against ***mf*TfR1**) were employed as secondary antibodies. Prior to imaging, nuclei were counterstained with DAPI. All sections were visualized using a confocal laser scanning microscope.

### Overexpression assay

To confirm transfection efficiency, FHM cells seeded in 6-well plates (60% confluency) were transfected with 4 µg of pEGFP-*mf*TfR1 plasmids. Post-transfection (48 h), cells were rinsed twice with ice-cold PBS, harvested using a cell scraper, and aliquoted into 3.5 mL tubes. Samples were bifurcated: one aliquot was stored in TRIzol for RNA extraction and subsequent qPCR, while the other was lysed in RIPA buffer supplemented with 1% PMSF for immunoblotting. To assess the impact of *mf*TfR1 on SDDV replication, transfected FHM cells were infected with SDDV (MOI = 5) 24 h post-transfection and harvested at intervals (0.5–120 hpi). Subsets of cells were processed as above for RNA and protein analysis. Remaining cells underwent three freeze-thaw cycles, and viral titers were quantified via the Reed-Muench method. IFA were conducted in 24-well plates containing glass coverslips (WHB, China). Cells were probed with monoclonal anti-MCP and polyclonal anti-*mf*TfR1 antibodies, followed by Alexa Fluor 555-labeled anti-mouse IgG and Alexa Fluor 488-labeled anti-rabbit IgG secondary antibodies for co-localization analysis.

### Cell viability assay

Cell viability was assessed using the CCK-8 assay kit (Solarbio, China), after the cells were treated with test drugs, 10 μL of CCK-8 solution was added to each well. The cells were then incubated for 1 h, followed by washing with PBS. The absorbance at OD_450_ was measured using a spectrophotometer. The cell viability was calculated by comparing the OD values with those of the control group.

### Prediction of *mf*TfR1 Interaction with SDDV Proteins

The domains of ***mf*TfR1** were predicted using <u>AlphaFold3</u> to generate the protein structure, which was further compared with the mTfR1 and hTfR1 domains using SnapGene software and Chimera X for structural analysis. The potential interactions between ***mf*TfR1** and various SDDV proteins were predicted using <u>AlphaFold3</u> (https://alphafoldserver.com/, followed by confirmation through Co-IP to validate the specific binding between ***mf*TfR1** and MCP. The binding site of ***mf*TfR1** with MCP was determined by <u>AlphaFold3</u> to obtain the protein structure, and the amino acids at the predicted binding site were analyzed using Chimera X.

### Antibody blocking

MFF-1 cells were seeded in 6-well and 24-well plates and treated with ***mf*TfR1** pAb at 0.16 μg/mL, 1.6 μg/mL, 16 μg/mL or with rabbit IgG as a control, for 2 h on ice. After washing with PBS, cells were collected for RNA extraction (TRIzol) or protein analysis (RIPA buffer with 1% PMSF). Cells were then incubated with SDDV (MOI = 5) at 4 °C for 1 h, washed, and cultured in antibody-containing medium at 27 °C. CPE was monitored for 3 dpi. Samples for qPCR (all antibody concentrations) and Western blot (16 μg/mL ***mf*TfR1** pAb) were collected at 0.5-120 hpi. Viral titers were determined by freeze-thawing cells three times and using the Reed and Muench method. For IFA, cells treated with 16 μg/mL ***mf*TfR1** pAb were fixed at 2dpi and 3dpi with 3% paraformaldehyde. MCP mAb was used as the primary antibody, Alexa Fluor 555-labeled donkey anti-mouse IgG as the secondary antibody, and nuclei were counterstained with Hoechst 33342.

### Ferristtin II treatment assay

Ferristatin II was purchased from GLPBIO (California, USA) and diluted in DMSO to final concentrations of 25 μM, 50 μM, and 100 μM. It was added to MFF-1 cell culture medium 5 h prior to SDDV infection, qPCR and Western blot were performed as described above.

### Coimmunoprecipitation (Co-IP) assay

To interrogate protein-protein interactions, Co-IP assays were performed as follows: FHM cells (60–70% confluency in 6-well plates) were transfected with plasmid combinations (4 µg/mL) via jetPRIME transfection reagent (Polyplus, France). At 48 h post-transfection, cells were rinsed twice with ice-cold PBS, homogenized in 100 µL PMSF-supplemented lysis buffer (1% v/v), and agitated on ice for 30 min. Lysates were centrifuged at 12,000 × g (4°C, 20 min) to remove debris. A 60 µL aliquot of the clarified supernatant was incubated with 20 µL anti-Flag or anti-GFP-conjugated magnetic beads (Beyotime, China) at 4°C for 2 h, with the residual lysate retained as input controls. Post-incubation, bead-bound complexes were magnetically separated, washed 6 times with lysis buffer (5 min per wash under gentle rotation), and resuspended in SDS-PAGE loading buffer. Protein eluates were denatured and subjected to immunoblotting analysis alongside input samples to validate specific interactions.

### Immunoprecipitation (IP)

Cells were plated in 6-well plates and subjected to infection with SDDV strain ZH-06/20 or mock treatment. At 1 hpi, cells underwent lysis on ice using 1 mL RIPA buffer supplemented with protease inhibitors. Lysates were clarified by centrifugation at 12,000 × g (4°C, 20 min). To minimize nonspecific interactions, supernatants were precleared with 2 μL anti-rabbit IgG and 20 μL protein A/G magnetic beads under gentle agitation at 4°C for 1 h. Following magnetic removal of preclearing beads, lysates were incubated overnight at 4°C with 1 μg of *mf*TfR1 polyclonal antibody (pAb) or Src pAb under continuous rotation. Subsequently, 20 μL protein A/G magnetic beads were added to each sample and incubated for 2 h at 4°C with gentle mixing. Beads bound to *mf*TfR1 or Src complexes were sequentially washed in PBS to remove unbound proteins, followed by elution for downstream analysis. ZH-06/20

### Peptide synthesis of *mf*TfR1-Binding sites using the pETB2M vector

Peptides corresponding to the ***mf*TfR1**-binding sites with >95% purity were synthesized by GeneCreate Biological Engineering Co., Ltd. (Wuhan, China). As a control, an empty pETB2M vector was used to synthesize unrelated peptides under the same conditions. All peptides were further purified using reverse-phase HPLC (RP-HPLC) to ensure > 95% purity, lyophilized, and stored at -80 °C. The purified peptides were quantified using a BCA Protein Assay Kit (Thermo Fisher Scientific, USA) following the manufacturer’s instructions. The synthesized peptides, including control peptides, were stored at −80 °C for subsequent experiments.

### Receptor blocking assay

ZH-06/20 was pre-incubated with the ***mf*TfR1**-binding site peptide at final concentrations of 100µg/mL in serum-free DMEM for 1 h at 4°C. Following pre-incubation, the peptide-treated SDDV was added to the MFF-1 cells at an MOI of 5 and incubated for 1 h at 27 °C. After virus adsorption, the cells were washed three times with PBS and further cultured in DMEM containing 5% FBS for 48 h. At 1-3 dpi, Cells were harvested for qPCR and IFA analysis of SDDV RNA levels. Mock groups included MFF-1 cells or ZH-06/20 incubated with peptides derived from the empty pETB2M vector under the same conditions.

### Viral binding assay

Cells were plated in 12-well plates and either transfected with target-specific siRNA for 48 h or pre-incubated with methyl-β-cyclodextrin (MβCD, Sigma) or Nystatin (Sigma, USA) for 1 h. To synchronize viral attachment, cells were chilled on ice for 20 min, followed by removal of culture medium and addition of 200 µL SDDV strain ZH-06/20 (MOI = 10) in prechilled buffer. After 1h incubation at 4°C, unbound virions were eliminated by three washes with ice-cold PBS. Cellular pellets were harvested for qPCR analysis to assess bound viral genome copies.

### Viral entry assay

MFF-1 cells cultured on 24-well plates were transfected with siRNA for 48 h prior to infection. Post-internalization (27°C), cells were fixed with 4% paraformaldehyde (15 min, room temperature). For intracellular staining, cells were permeabilized with 0.1% Triton X-100 (10 min) and blocked with 5% bovine serum albumin (BSA) to minimize nonspecific binding. Both permeabilized and non-permeabilized cells were probed overnight at 4°C with MCP mAb, washed, and incubated with Alexa Fluor 555-conjugated donkey anti-mouse IgG (1 h). Fluorescence signals from ≥35 cells per condition were quantified using ImageJ software.

### Statistical analysis

The data are expressed as means ± standard deviation (SD) based on three independent experiments. Statistical analysis was carried out using SPSS 16.0. Comparisons between control and experimental groups were conducted using Student’s t-test and one-way Analysis of Variance (ANOVA). Statistical significance was considered when 0.01 < \**p* < 0.05, \*\**p* < 0.01, and non-significant differences were marked as “ns.” Groups were labeled with different letters (a, b, c, d) to denote significant differences.

## ACKNOWLEDGMENTS

This work was funded by National Natural Science Foundation of China under No. 32403075; the key areas R&D Program of Guangdong Province under No. 2021B0202040002; Guangdong Basic and Applied Basic Research Foundation under No. 2023A1515110938; and China Postdoctoral Science Foundation under No. 2024M753721.

## Data Availability Statement

The authors confirm that the data supporting the findings of this study are available within the article [and/or] its supplementary materials.

